# Mutations in two SARS-CoV-2 variants of concern reflect two distinct strategies of antibody escape

**DOI:** 10.1101/2021.07.23.453327

**Authors:** Sebastian Fiedler, Viola Denninger, Alexey S. Morgunov, Alison Ilsley, Roland Worth, Georg Meisl, Catherine K. Xu, Monika A. Piziorska, Francesco Ricci, Anisa Y. Malik, Sean R. A. Devenish, Matthias M. Schneider, Vasilis Kosmoliaptsis, Adriano Aguzzi, Akiko Iwasaki, Heike Fiegler, Tuomas P. J. Knowles

## Abstract

Understanding the factors that contribute to antibody escape of SARS-CoV-2 and its variants is key for the development of drugs and vaccines that provide broad protection against a variety of virus variants. Using microfluidic diffusional sizing, we determined the dissociation constant (*K*_D_) for the interaction between receptor binding domains (RBDs) of SARS-CoV-2 in its original version (WT) as well as alpha and beta variants with the host-cell receptor angiotensin converting enzyme 2 (ACE2). For RBD-alpha, the ACE2-binding affinity was increased by a factor of ten when compared with RBD-WT, while ACE2-binding of RBD-beta was largely unaffected. However, when challenged with a neutralizing antibody that binds to both RBD-WT and RBD-alpha with low nanomolar *K*_D_ values, RBD-beta displayed no binding, suggesting a substantial epitope change. In SARS-CoV-2 convalescent sera, RBD-binding antibodies showed low nanomolar affinities to both wild-type and variant RBD proteins—strikingly, the concentration of antibodies binding to RBD-beta was half that of RBD-WT and RBD-alpha, again indicating considerable epitope changes in the beta variant. Our data therefore suggests that one factor contributing to the higher transmissibility and antibody evasion of SARS-CoV-2 alpha and beta is a larger fraction of viruses that can form a complex with ACE2. However, the two variants employ different mechanisms to achieve this goal. While SARS-CoV-2 alpha RBD binds with greater affinity to ACE2 and is thus more difficult to displace from the receptor by neutralizing antibodies, RBD-beta is less accessible to antibodies due to epitope changes which increases the chances of ACE2-binding and infection.

## Introduction

The spread of SARS-CoV-2 is governed by a combination of efficient transmission of the virus, and its ability to resist virus neutralization. Therefore, to successfully fight the SARS-CoV-2 pandemic it is crucial to fully understand the receptor recognition mechanisms, as this interaction regulates transmission, pathogenesis and host range^1^.

The hallmark of the adaptive immune response towards SARS-CoV-2 resides in the generation of neutralizing antibodies (NAbs) that prevent the virus from binding to the angiotensin-converting enzyme 2 (ACE2) receptor on the host cell. Cell entry is driven by the SARS-CoV-2 spike protein, a trimeric transmembrane glycoprotein comprising S1 and S2 subunits, with the S1 subunit mediating viral binding to ACE2^2^. Given that the spike protein has become a prime target for vaccines, the development of mutations within the antigenic sites of the S1 receptor binding domain (RBD) raises significant concerns of viral escape of humoral immunity.

The rapid global spread of SARS-CoV-2 has seen significant changes in viral fitness, and more than 4,000 variants have been reported to date^3^. The emergence of B.1.1.7 (SARS-CoV-2 alpha) and B.1.351 (SARS-CoV-2 beta) SARS-CoV-2 variants in the United Kingdom and South Africa, respectively, have raised specific concerns given that mutations in their respective spike-protein sequences have not only led to an increase in transmission, but both mutant strains have also become impervious to multiple NAbs that target both the RBD and N-terminal domain (NTD) of the original spike protein^4^.

The alpha and beta variants share a N501Y mutation which is of particular importance considering that amino acid N501 is thought to be instrumental in stabilizing the interaction between the viral spike protein and the ACE2 receptor^1^. With the N501Y mutation, the altered interface seems to be accompanied by a change in the interaction network, with an increase in the number of hydrogen bonds and van der Waals contacts and leading to an increase in the binding affinity between the viral RBD region and the ACE2 receptor^5,6^. Interestingly, the N501Y mutation only showed a modest impairment on the efficacy of late-stage clinical trial monoclonal antibodies (mAbs) REGN10933 and REGN10987 (Regeneron) and AZD1061 and AZD8895 (AstraZeneca)^7^. Furthermore, pseudoviruses with the N501Y mutation exhibited a similar neutralization potency to wild-type SARS-CoV-2 when exposed to vaccinated and convalescent serum samples^4^. Although the neutralization potential of these currently administered mAbs and vaccines is unaffected by SARS-CoV-2 alpha, not all mAbs raised against the wild type are effective in neutralizing this variant. In the structures of 38 antibody fragments (Fabs) in complex with RBD, the complementarity-determining regions 1 for 47% of these Fabs were in direct contact with N501, and as a result the neutralization efficacy of various mAbs were affected by this mutation^8^.

In contrast to the alpha variant, the RBD of the beta variant has additional K417N and E484K mutations, raising specific concerns regarding viral antibody escape. This is supported by observations of pseudoviruses with a triple spike mutation (N501Y, K417N, E484K) which either abrogated or significantly resisted neutralization in convalescent and post-vaccinated sera^8,9^. Interestingly, pseudoviruses carrying only single E484K, K417N or double K417N/E484K mutations showed only two-fold increases in transmission rates relative to wild-type pseudoviruses^4^. Yet, when combined with N501Y these mutations exhibited higher transmission rates, with N501Y/E484K showing a 13-fold increase relative to wild-type pseudoviruses. Interestingly, E484K and N501Y/E484K pseudoviruses partly resisted neutralization by post-vaccinated sera, but pseudoviruses with the K417N/E484K or the triple mutation exhibited the highest resistance to neutralization^4^. The ramification of these variants was highlighted in a trial which revealed that the double dose of AZD1222 vaccine only had a 10.4% efficacy at preventing mild to moderate symptoms induced by the beta variant^9^.

In this study, we have used RBD-WT, RBD-alpha, and RBD-beta to quantify the impact of various RBD mutations (N501Y, K417N and E484K) on the binding affinity to ACE2 and a monoclonal Nab raised against the original SARS-CoV-2 (Wuhan-Hu-1 sequence). Then, we evaluated the same wild-type and variant RBDs in microfluidic antibody affinity profiling (MAAP) to measure affinity and concentrations of anti-RBD antibodies in convalescent serum obtained from individuals that recovered from an infection with the original SARS-CoV-2 variant. Unlike measurement of titers, determination of both affinity and concentration allows us to distinguish changes in affinity across the entire antibody population from loss of binding of a subset of antibodies. Correlating these serum-antibody affinities and concentrations with their ability to disrupt ACE2/spike S1 complexes enabled us to quantify the response of anti-RBD-WT antibodies in serum when challenged with RBD-alpha or RBD-beta. Our results suggest that while the anti-RBD-WT antibodies are effective against RBD-WT, they are less effective against both RBD-alpha and RBD-beta variants. More importantly, RBD-alpha and RBD-beta follow different approaches to evade the immune response. For RBD-alpha, antibody escape is mainly driven by an increase in the binding affinity to the ACE2 receptor, while RBD-beta seems to prevent antibody binding by changing key epitopes on the surface of the protein. As a result, both variants are more effective in binding to ACE2 on the surface of host-cells thus increasing their chance of successful cell-entry

## Results and Discussion

Antibody escape by SARS-CoV-2 variants can be achieved by different mechanisms including an increased binding affinity to ACE2 or through the alteration of key NAb binding epitopes on the spike surface. To deconvolute these two escape mechanisms, we measured the ACE2 binding affinity of RBD-WT, RBD-alpha, RBD-beta, RBD-K417N, and RBD-E484K (**Figure 1A, Table 1**) and then investigated binding affinities of the same RBDs to a monoclonal NAb that was obtained from a patient infected with the original SARS-CoV-2 variant (Wuhan-Hu-1 sequence; **Figure 1B, Table 1**). The *K*_D_ of ACE2 binding to RBD-WT was 46 nM (CI 95%: 28–74), which is in line with previously observed values^10–15^. RBD-alpha showed a considerably higher ACE2-binding affinity than RBD-WT with *K*_D_ being reduced approximately by a factor of 10. For RBD-beta and the single mutant RBD-K417N, the ACE2-affinties were within experimental error of the RBD-WT *K*_D_. However, E484K binding to ACE2 has a *K*_D_ that is approximately 8 times higher than for RBD-WT resulting in weaker binding affinity. The triple mutant RBD-beta therefore combines mutations that promote ACE2 binding (N501Y) with some that either have a limited effect (K417N) or even impede ACE2 binding (E484K), rationalizing the fact that the overall binding affinity of RBD-beta is comparable to the WT. Thus, an increase in ACE2 binding affinity is unlikely to be the main mechanism of immune escape for SARS-CoV 2 beta and it must draw additional advantages from carrying these mutations.

**Figure 1.**
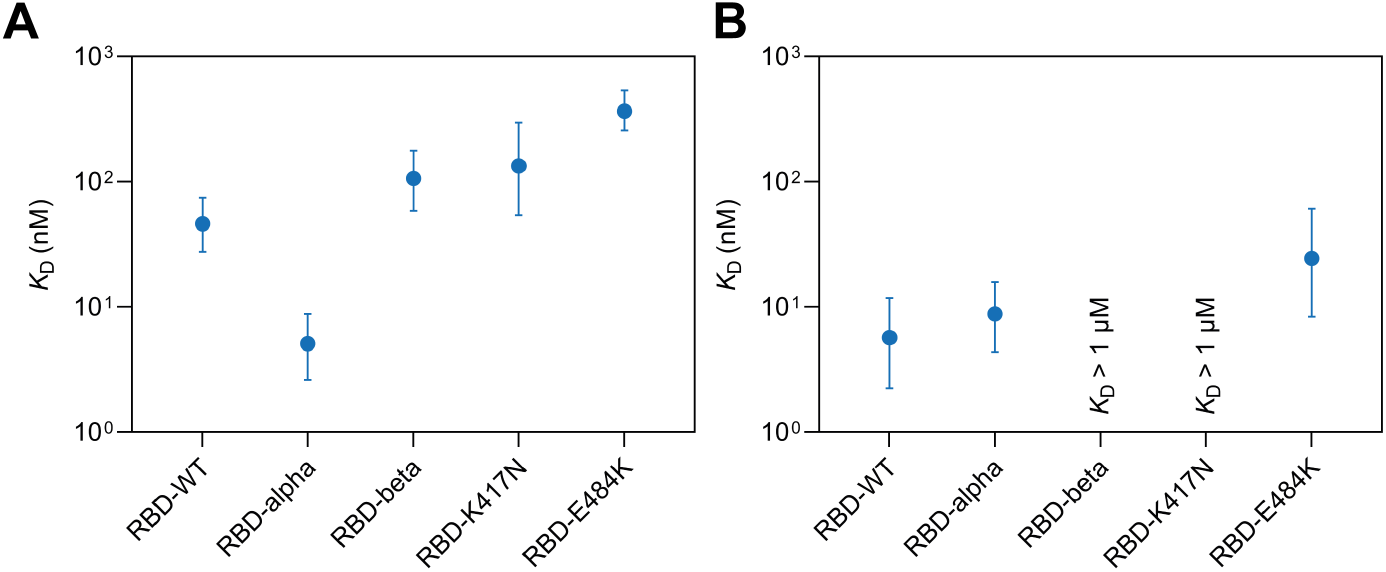
Dissociation constants, *K*_D_, of different variants of SARS-CoV-2 RBD binding to (A) ACE2 and (B) a neutralizing monoclonal antibody. Equilibrium binding was measured by microfluidic diffusional sizing for various concentrations of fluorescently labeled RBD variants and unlabeled ACE2 or unlabeled NAb. *K*_D_ values were determined from modes of posterior probability distributions obtained by Bayesian inference^18,19^. Error bars are 95% credible intervals. Refer to Table 1 for nomenclature of RBD proteins.

**Table 1.** Summary of recombinant proteins.

| SARS-CoV-2 proteins |  |  |  |  |  |  |
| --- | --- | --- | --- | --- | --- | --- |
| Protein name | SARS-CoV-2 variant | Pango lineage | Vendor | Article number | Amino-acid sequence | Mutated amino-acid residues |
| RBD-WT | Original | A | Sino Biological | 40592-V08H | R319-F541 |  |
| RBD-alpha | Alpha | B.1.1.7 | Sino Biological | 40592-V02H1 | R319-F541 | N501Y |
| RBD-beta | Beta | B.1.351 | Sino Biological | 40592-V08H85 | R319-F541 | K417N, E484K, N501Y |
| RBD-K417N |  |  | Sino Biological | 40592-V08H59 | R319-F541 | K417N |
| RBD-E484K |  |  | Sino Biological | 40592-V08H84 | R319-F541 | E484K |
| S1-WT | Original | A | Acro Biosystems | S1N-C52H4 | V16-R685 |  |
| S1-alpha | Alpha | B.1.1.7 | Acro Biosystems | S1N-C52Hg-100ug | V16-R685 | N501Y |
| S1-beta | Beta | B.1.351 | Sino Biological | 40591-V08H10 | M1-R685 | K417N, E484K, N501Y, D614G |
| Host-cell protein |  |  |  |  |  |  |
| Protein name | N/A | N/A | Vendor | Article number | Amino-acid sequence | N/A |
| ACE2 |  |  | Acro Biosystems | AC2-H52H8 | Q18-S740 |  |
| Recombinantly produced neutralizing monoclonal antibody |  |  |  |  |  |  |
| Protein name | N/A | N/A | Vendor | Article number | N/A | N/A |
| NAb |  |  | Acro Biosystems | SAD-S35 |  |  |

To investigate the effect of the alpha and beta variant mutations on antibody binding, we used MDS to measure affinities to a monoclonal NAb that was obtained from a patient infected with the original SARS-CoV-2 variant (**Figure 1B**). RBD-WT and RBD-alpha displayed similar *K*_D_ values (5.7 nM (CI 95%: 2.2–11.8) and 8.8 nM (CI 95%: 4.4–15.8), respectively). For RBD-E484K, *K*_D_ increased by a factor of 4 as compared with RBD-WT, resulting in a considerably reduced binding affinity to this antibody. Remarkably, the Nab did not show any binding to RBD-K417N up to the maximum tested concentration of 1 µM. Consequently, the triple mutant RBD-beta which contains K417N was also not bound. This suggests that RBD-beta mutations K417N and E484K do not increase the binding affinity to ACE2 but rather cause some antibodies to lose their ability to bind to the RBD domain.

This was further confirmed by a recent study that combined kinetic binding experiments by surface plasmon resonance (SPR) with structural data obtained by cryo-electron microscopy ^16^. In this study, monoclonal antibodies targeting epitopic region RBD2^17^ did not bind to RBD-beta, while antibody affinity to RBD-alpha was found to be similar to RBD-WT^16^. In addition, the ACE2 binding affinity was increased for RBD-alpha while there was no difference between RBD-beta and RBD-WT^16^. The structural data also suggested that N501Y increases ACE2 affinity due to a hydrophobic interaction between Y501 and Y41 in ACE2, accompanied by cation-π interactions with K353 in ACE2^16^. In contrast, the RBD-beta– ACE2 complex lacks salt bridges between RBD-beta K417 and ACE2 D30 and between RBD-beta E484 and ACE2 K31, which seems to counteract the increased ACE2 affinity mediated by N501Y^16^.

Next, we investigated how our observations on recombinant proteins translate to three convalescent serum samples from individuals who had been infected with the original SARS-CoV-2 variant. We used microfluidic antibody affinity profiling (MAAP)^18–20^ followed by an in-solution receptor-binding competition assay^19,20^ using RBD-WT, RBD-alpha and RBD-beta. While MAAP quantifies the affinity and the concentration of the polyclonal antibody mixture against RBD, the in-solution receptor-binding competition assay identifies NAbs that can displace spike S1 from the ACE2 receptor.

In all three serum samples, antibodies bound tightly to each of the three different RBDs with *K*_D_ in a range of 1–4 nM (**Figure 2A**). The concentration of antibody binding sites varied over one order of magnitude from 10 nM to 100 nM. Thus, all three serum samples contained antibodies that were able to bind to all three variants of RBD with high affinity. Overall, sample 3973 contained the highest concentration of antibodies binding to all three types of RBD, followed by sample 3541 and sample 3707. Similar concentrations of binding antibodies against RBD-WT and RBD-alpha were found in each of the three serum samples, whereas the concentration of antibodies that bind to RBD-beta were lower than the other RBDs in two samples and higher in one sample, again highlighting that there may be significant differences in the binding epitope for RBD-beta.

**Figure 2.**
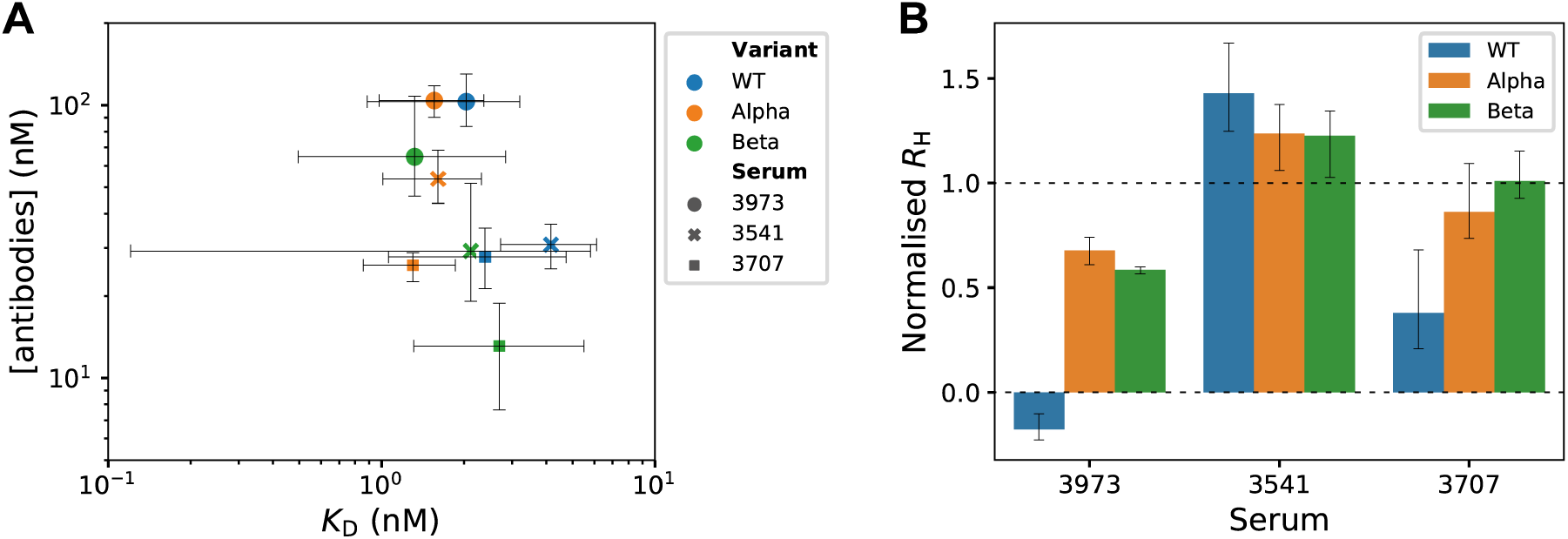
Microfluidic antibody-affinity profiling and in-solution receptor-binding competition assay in SARS-CoV-2 convales-cent serum. (A) Equilibrium binding was measured by microfluidic diffusional sizing for various concentrations of fluorescently labeled RBD variants and dilutions of convalescent sera. *K*_D_ values and antibody concentrations were determined from modes of posterior probability distributions obtained by Bayesian inference^18,19^. Error bars are 95% credible intervals. (B) In-solution receptor-binding competition assay identifies polyclonal antibodies that displace spike S1 from the ACE2 receptor. At normalized *R*_h_ = 0, all S1 was displaced from ACE2, and the measured *R*_h_ is the same as for free ACE2. At normalized *R*_h_ = 1, S1 was not displaced from ACE2 and the measured *R*_h_ is the same as for ACE2/S1 complex. Error bars are standard deviations obtained from triplicate measurements.

Through an in-solution receptor-binding competition assay^20^ (**Figure 2B**), we found that the antibodies in sample 3541 were unable to displace any of the three different S1 domains (WT, alpha, beta) from the ACE2 receptor, which suggests that the majority of antibodies in this sample do not target the receptor binding motif of the RBD and are thus unable to outcompete ACE2. Out of the other two serum samples, 3973 showed slightly increased displacement for all types of S1, an effect which is likely caused by the overall higher antibody concentration in sample 3973 (**Figure 2A**). Interestingly however, samples 3973 and 3707 displayed the strongest displacement efficacy for S1-WT while both S1-alpha and S1-beta showed reduced displacement from ACE2 in line with increased transmission of the variants.

Based on these observations, we argue that for RBD-alpha, the reduced ability of serum antibodies to outcompete ACE2 binding to RBD can be explained by the increased affinity of the RBD for the ACE2 receptor (**Figure 1A**). Quantitatively, the inhibition of spike/ACE2 binding by NAbs is described by a ternary equilibrium in which antibodies compete with ACE2 for the binding to spike^20^. The parameters that govern this ternary equilibrium are the *K*_D_ of spike/ACE2 binding, the *K*_D_ of antibody/spike binding and the total concentrations of spike, ACE2, and antibodies. Our data suggest that the affinity and the concentration of antibodies against both RBD-WT and RBD-alpha in convalescent serum are largely similar (**Figure 2A**). Consequently, amongst the parameters tested, for RBD-alpha only the affinity of RBD to ACE2 was increased, leading to the formation of a higher proportion of RBD/ACE2 complexes as compared with RBD-WT. During a SARS-CoV-2 infection, re-infection, or infection of vaccinated individuals, the same ternary equilibrium composed of serum antibodies, spike protein and ACE2 governs the amount of virus that binds to host cells. Thus, with the same level of anti-RBD antibodies and the same number of virus particles, SARS-CoV-2 alpha can attach to more ACE2 receptors (i.e., host cells) and therefore be more transmissible than the WT variant.

By contrast, the reduced displacement efficacy for RBD-beta can be explained by a lower number of serum antibodies that are able to bind to RBD-beta as compared with WT. We have found a reduced concentration of antibodies binding to RBD-beta in serum samples 3973 and 3707 in comparison with RBD-WT and RBD-alpha. In contrast to RBD-alpha, both the affinity for ACE2 as well as the affinity of the polyclonal antibody response was largely unaffected for RBD-beta. In terms of the ternary equilibrium of spike/ACE2 and antibodies, a reduction in anti-RBD antibody concentration again leads to a higher proportion of RBD/ACE2 complexes and higher occupancies of host-cell receptors and, thus, might be related to increased transmission.

## Conclusion

Our data provides insights into the different strategies adopted by the SARS-CoV-2 alpha and beta variants to evade antibodies generated during an earlier infection with SARS-CoV-2. We argue that one of the mechanisms by which SARS-CoV-2 can boost its transmission is by increasing the fraction of virus particles that occupy the surface of host cells. This can be achieved either through higher-affinity binding to the host-cell receptor or through mutations in antibody binding epitopes on the surface of the spike protein in order to prevent antibody binding. Our data provide quantitative evidence that the SARS-CoV-2 alpha and beta variants adopt different strategies to achieve higher occupancy on the surface of host cells. RBD-alpha shows a ten times higher binding affinity to ACE2 than RBD-WT with little effect on affinities and concentrations of binding antibodies contained in SARS-CoV-2 convalescent serum. Thus, to boost its transmission, SARS-CoV-2 alpha seems to take the approach of increasing its binding affinity to ACE2. On the other hand, the two additional mutations (K417N and E484K) in SARS-CoV-2 beta do not seem to be beneficial for ACE2 binding, instead reducing its affinity back to the levels seen for WT. However, in SARS-CoV-2 convalescent serum, the concentration of antibodies able to bind RBD beta is two-fold lower than that of antibodies binding RBD WT. Thus, during re-infection, SARS-CoV-2 beta potentially evades the initial antibody response by interfering directly with antibody binding rather than by an increased binding affinity to ACE2.

Our data provides quantitative insights on the molecular mechanisms of antibody escape employed by SARS-CoV-2 variants of concern. Such quantitative data are essential to understand how new virus variants evolve and can pave the way towards the development of broad therapeutics and vaccines that are not only effective against a wide range of existing viral variants but also cover the development of any future versions of the virus that may evolve.

## Materials and Methods

### Fluorescent labeling of proteins

Recombinant proteins were labeled with Alexa Fluor™ 647 NHS ester (Thermo Fisher) as described previously.^20^ In brief, 50 µg of protein solution was mixed with dye at a three-fold molar excess in the presence of NaHCO_3_ (Merck) buffer at pH 8.3, incubated at 4 °C overnight, and unbound label was removed by size-exclusion chromatography (SEC) on an ÄKTA pure system (Cytiva). SEC runs of RBD proteins were performed on a Superdex 75 Increase 10/300 column (Cytiva), while S1 and ACE2 were run on a Superdex 200 Increase 3.2/300 column (Cytiva). Labeled and purified proteins were stored at −80 °C in PBS pH 7.4 containing 10% (w/v) glycerol as cryoprotectant.

### Equilibrium affinity measurements by microfluidic diffusional sizing

Binding affinity of RBD–ACE2 and RBD–NAb was measured on a Fluidity One-W Serum (Fluidic Analytics) as described previously.^19,20^ In brief, various concentrations of fluorescently labeled RBD and unlabeled ACE2 or NAb were mixed and incubated on ice for 1 h. The change in hydrodynamic radius (*R*_h_) upon binding was measured in duplicate. *K*_D_ values were determined by Bayesian inference as described previously.^18,19^

### Origin of serum samples

Anti-SARS-CoV-2 seropositive human serum samples (convalescent) were obtained from BioIVT. BioIVT sought informed consent from each subject, or the subjects legally authorized representative, and appropriately documented this in writing. All samples are collected under IRB-approved protocols.

### Microfluidic antibody-affinity profiling (MAAP)

MAAP on a Fluidity One-W Serum was used to determine concentration of antibody-binding sites and *K*_D_ of antibodies against RBD in serum samples of SARS-CoV-2 seropositive individuals as described previously.^19,20^ In brief, various concentrations of fluorescently labeled RBD and convalescent serum were mixed and incubated on ice for 1 h. The change in *R*_h_ upon binding was measured in duplicate. Concentration of antibody binding sites and *K*_D_ values were determined by Bayesian inference as described previously.^18,19^

### Serum-antibody mediated displacement of ACE2 from S1 by using microfluidic diffusional sizing

Displacement of ACE2 from SARS-CoV-2 S1 was measured on a Fluidity One-W Serum as described previously.^19,20^ S1 is used to improve S/N as S1 binding induces a larger size change than RBD upon binding to ACE2. In brief, labeled ACE2 (10 nM) and unlabeled S1 (30 nM) were mixed with convalescent serum (90%) and incubated on ice for 30 min. The change in *R*_h_ upon displacement of S1 was measured in triplicate. The *R*_h_ of unbound labeled ACE2 at a concentration of 10 nM and ACE2–S1 mix (10 Nm and 30 nM, respectively) in the absence of convalescent serum were used as reference values for 100% displacement and 0% displacement, respectively.

## Notes

The authors declare the following competing financial interest(s): T.P.J.K. is a member of the board of directors of Fluidic Analytics. A.A. is a member of the scientific advisory committee of Fluidic Analytics. V.K., M.M.S., and G.K., C.K.X. are consultants for Fluidic Analytics. S.F., V.D., A.S.M., A.I., R.W., M.A.P., F. R., A.Y.M., S.R.A.D., and H.F. are employees of Fluidic Analytics.

## Author Contributions

Study design: S.F., A.S.M., H.F. Experiments: S.F., V.D., A.I., M.A.P., F.R., A.Y.M. Data Analysis: S.F., A.S.M., G.M., C.K.X. Study advice: S.R.A.D., M.M.S., A.A., V.K., A.I. Study supervision: H.F., T.P.J.K. Manuscript writing: S.F., R.W., H.F.

## Acknowledgements

The authors thank the entire team at Fluidic Analytics for doing things that make a real difference every day. A.A. was funded by institutional core funding by the University of Zurich and the University Hospital of Zurich, Driver Grant 2017DRI17 of the Swiss Personalized Health Network (SPHN), Distinguished Scientist Award of the NOMIS Foundation, a Grant of the European Research Council (ERC Prion2020 670958), and an Innovation Fund of the University Hospital Zurich. V.K. was funded by an NIHR fellowship (PD-2016-09-065) and acknowledges support as a PI Terasaki Scholar. A.I. is an investigator of the Howard Hughes Medical Institute. T.P.J.K. is grateful for financial support by the Biotechnology and Biological Sciences Research Council and the European Research Council.

## References

(1) Shang, J.; Ye, G.; Shi, K.; Wan, Y.; Luo, C.; Aihara, H.; Geng, Q.; Auerbach, A.; Li, F. Structural Basis of Receptor Recognition by SARS-CoV-2. Nature 2020, 581 (7807), 221–224. https://doi.org/10.1038/s41586-020-2179-y.

(2) Huang, Y.; Yang, C.; Xu, X. feng; Xu, W.; Liu, S. wen. Structural and Functional Properties of SARS-CoV-2 Spike Protein: Potential Antivirus Drug Development for COVID-19. Acta Pharmacologica Sinica. Springer US 2020, pp 1141–1149. https://doi.org/10.1038/s41401-020-0485-4.

(3) Bian, L.; Gao, F.; Zhang, J.; He, Q.; Mao, Q.; Xu, M.; Liang, Z. Effects of SARS-CoV-2 Variants on Vaccine Efficacy and Response Strategies. Expert Review of Vaccines. Taylor & Francis 2021, pp 1–9. https://doi.org/10.1080/14760584.2021.1903879.

(4) Kuzmina, A.; Khalaila, Y.; Voloshin, O.; Keren-Naus, A.; Boehm-Cohen, L.; Raviv, Y.; Shemer-Avni, Y.; Rosenberg, E.; Taube, R. SARS-CoV-2 Spike Variants Exhibit Differential Infectivity and Neutralization Resistance to Convalescent or Post-Vaccination Sera. Cell Host Microbe 2021, 29 (4), 522-528.e2. https://doi.org/10.1016/j.chom.2021.03.008.

(5) Starr, T. N.; Greaney, A. J.; Hilton, S. K.; Ellis, D.; Crawford, K. H. D.; Dingens, A. S.; Navarro, M. J.; Bowen, J. E.; Tortorici, M. A.; Walls, A. C.; King, N. P.; Veesler, D.; Bloom, J. D. Deep Mutational Scanning of SARS-CoV-2 Receptor Binding Domain Reveals Constraints on Folding and ACE2 Binding. Cell 2020, 182 (5), 1295-1310.e20. https://doi.org/10.1016/j.cell.2020.08.012.

(6) Khan, A.; Zia, T.; Suleman, M.; Khan, T.; Ali, S. S.; Abbasi, A. A.; Mohammad, A.; Wei, D. Q. Higher Infectivity of the SARS-CoV-2 New Variants Is Associated with K417N/T, E484K, and N501Y Mutants: An Insight from Structural Data. J. Cell. Physiol. 2021, No. January, 1–13. https://doi.org/10.1002/jcp.30367.

(7) Wang, P.; Liu, L.; Iketani, S.; Luo, Y.; Guo, Y.; Wang, M.; Yu, J.; Zhang, B.; Kwong, P. D.; Graham, B. S.; Mascola, J. R.; Chang, J. Y.; Yin, M. T.; Sobieszczyk, M.; Kyratsous, C. A.; Shapiro, L.; Sheng, Z.; Nair, M. S.; Huang, Y.; Ho, D. D. Increased Resistance of SARS-CoV-2 Variants B.1.351 and B.1.1.7 to Antibody Neutralization. bioRxiv Prepr. Serv. Biol. 2021. https://doi.org/10.1101/2021.01.25.428137.

(8) Supasa, P.; Zhou, D.; Dejnirattisai, W.; Liu, C.; Mentzer, A. J.; Ginn, H. M.; Zhao, Y.; Duyvesteyn, H. M. E.; Nutalai, R.; Tuekprakhon, A.; Wang, B.; Paesen, G. C.; Slon-Campos, J.; López-Camacho, C.; Hallis, B.; Coombes, N.; Bewley, K. R.; Charlton, S.; Walter, T. S.; Barnes, E.; Dunachie, S. J.; Skelly, D.; Lumley, S. F.; Baker, N.; Shaik, I.; Humphries, H. E.; Godwin, K.; Gent, N.; Sienkiewicz, A.; Dold, C.; Levin, R.; Dong, T.; Pollard, A. J.; Knight, J. C.; Klenerman, P.; Crook, D.; Lambe, T.; Clutterbuck, E.; Bibi, S.; Flaxman, A.; Bittaye, M.; Belij-Rammerstorfer, S.; Gilbert, S.; Hall, D. R.; Williams, M. A.; Paterson, N. G.; James, W.; Carroll, M. W.; Fry, E. E.; Mongkolsapaya, J.; Ren, J.; Stuart, D. I.; Screaton, G. R. Reduced Neutralization of SARS-CoV-2 B.1.1.7 Variant by Convalescent and Vaccine Sera. Cell 2021, 184 (8), 2201-2211.e7. https://doi.org/10.1016/j.cell.2021.02.033.

(9) Madhi, S. A.; Baillie, V.; Cutland, C. L.; Voysey, M.; Koen, A. L.; Fairlie, L.; Padayachee, S. D.; Dheda, K. Barnabas, S. L.; Bhorat, Q. E.; Briner, C.; Kwatra, G.; Ahmed, K.; Aley, P.; Bhikha, S.; Bhiman, J. N.; Bhorat, A. E.; du Plessis, J.; Esmail, A.; Groenewald, M.; Horne, E.; Hwa, S.-H.; Jose, A.; Lambe, T.; Laubscher, M.; Malahleha, M.; Masenya, M.; Masilela, M.; McKenzie, S.; Molapo, K.; Moultrie, A.; Oelofse, S.; Patel, F.; Pillay, S.; Rhead, S.; Rodel, H.; Rossouw, L.; Taoushanis, C.; Tegally, H.; Thombrayil, A.; van Eck, S.; Wibmer, C. K.; Durham, N. M.; Kelly, E. J.; Villafana, T. L.; Gilbert, S.; Pollard, A. J.; de Oliveira, T.; Moore, P. L.; Sigal, A.; Izu, A. Efficacy of the ChAdOx1 NCoV-19 Covid-19 Vaccine against the B.1.351 Variant. N. Engl. J. Med. 2021, 384 (20), 1885–1898. https://doi.org/10.1056/nejmoa2102214.

(10) Wrapp, D.; Wang, N.; Corbett, K. S.; Goldsmith, J. A.; Hsieh, C. L.; Abiona, O.; Graham, B. S.; McLellan, J. S. Cryo-EM Structure of the 2019-NCoV Spike in the Prefusion Conformation. Science (80-.). 2020 367 (6483), 1260–1263. https://doi.org/10.1126/science.aax0902.

(11) Shang, J.; Ye, G.; Shi, K.; Wan, Y.; Luo, C.; Aihara, H.; Geng, Q.; Auerbach, A.; Li, F. Structural Basis of Receptor Recognition by SARS-CoV-2. Nature 2020, 581 (7807), 221–224. https://doi.org/10.1038/s41586-020-2179-y.

(12) Lui, I.; Zhou, X.; Lim, S.; Elledge, S.; Solomon, P.; Rettko, N.; Zha, B. S.; Kirkemo, L.; Gramespacher, J.; Liu, J.; Muecksch, F.; Lorenzi, J. C. C.; Schmidt, F.; Weisblum, Y.; Robbiani, D.; Nussenzweig, M.; Hatziioannou, T.; Bieniasz, P.; Rosenburg, O.; Leung, K.; Wells, J. Trimeric SARS-CoV-2 Spike Interacts with Dimeric ACE2 with Limited Intra-Spike Avidity. bioRxiv Prepr. Serv. Biol. 2020. https://doi.org/10.1101/2020.05.21.109157.

(13) Glasgow, A.; Glasgow, J.; Limonta, D.; Solomon, P.; Lui, I.; Zhang, Y.; Nix, M. A.; Rettko, N. J.; Zha, S.; Yamin, R.; Kao, K.; Rosenberg, O. S.; Ravetch, J. V.; Wiita, A. P.; Leung, K. K.; Lim, S. A.; Zhou, X. X.; Hobman, T. C.; Kortemme, T.; Wells, J. A. Engineered ACE2 Receptor Traps Potently Neutralize SARS-CoV-2. Proc. Natl. Acad. Sci. U. S. A. 2020, 117 (45), 28046–28055. https://doi.org/10.1073/pnas.2016093117.

(14) Wang, Q.; Zhang, Y.; Wu, L.; Niu, S.; Song, C.; Zhang, Z.; Lu, G.; Qiao, C.; Hu, Y.; Yuen, K. Y.; Wang, Q.; Zhou, H.; Yan, J.; Qi, J. Structural and Functional Basis of SARS-CoV-2 Entry by Using Human ACE2. Cell 2020, 181 (4), 894-904.e9. https://doi.org/10.1016/j.cell.2020.03.045.

(15) Thomson, E. C.; Rosen, L. E.; Shepherd, J. G.; Spreafico, R.; da Silva Filipe, A.; Wojcechowskyj, J. A.; Davis, C.; Piccoli, L.; Pascall, D. J.; Dillen, J.; Lytras, S.; Czudnochowski, N.; Shah, R.; Meury, M.; Jesudason, N.; De Marco, A.; Li, K.; Bassi, J.; O’Toole, A.; Pinto, D.; Colquhoun, R. M.; Culap, K.; Jackson, B.; Zatta, F.; Rambaut, A.; Jaconi, S.; Sreenu, V. B.; Nix, J.; Zhang, I.; Jarrett, R. F.; Glass, W. G.; Beltramello, M.; Nomikou, K.; Pizzuto, M.; Tong, L.; Cameroni, E.; Croll, T. I.; Johnson, N.; Di Iulio, J.; Wickenhagen, A.; Ceschi, A.; Harbison, A. M.; Mair, D.; Ferrari, P.; Smollett, K.; Sallusto, F.; Carmichael, S.; Garzoni, C.; Nichols, J.; Galli, M.; Hughes, J.; Riva, A.; Ho, A.; Schiuma, M.; Semple, M. G.; Openshaw, P. J. M.; Fadda, E.; Baillie, J. K.; Chodera, J. D.; Rihn, S. J.; Lycett, S. J.; Virgin, H. W.; Telenti, A.; Corti, D.; Robertson, D. L.; Snell, G. Circulating SARS-CoV-2 Spike N439K Variants Maintain Fitness While Evading Antibody-Mediated Immunity. Cell 2021, 184 (5), 1171-1187.e20. https://doi.org/10.1016/j.cell.2021.01.037.

(16) Cai, Y.; Zhang, J.; Xiao, T.; Lavine, C. L.; Rawson, S.; Peng, H.; Zhu, H.; Anand, K.; Tong, P.; Gautam, A.; Lu, S.; Sterling, S. M.; Walsh, R. M.; Rits-Volloch, S.; Lu, J.; Wesemann, D. R.; Yang, W.; Seaman, M. S.; Chen, B. Structural Basis for Enhanced Infectivity and Immune Evasion of SARS-CoV-2 Variants. Science (80-.). 2021, eabi9745. https://doi.org/10.1126/SCIENCE.ABI9745.

(17) Tong, P.; Gautam, A.; Windsor, I.; Travers, M.; Chen, Y.; Garcia, N.; Whiteman, N. B.; McKay, L. G. A.; Lelis, F. J. N.; Habibi, S.; Cai, Y.; Rennick, L. J.; Duprex, W. P.; McCarthy, K. R.; Lavine, C. L.; Zuo, T.; Lin, J.; Zuiani, A.; Feldman, J.; MacDonald, E. A.; Hauser, B. M.; Griffths, A.; Seaman, M. S.; Schmidt, A. G.; Chen, B.; Neuberg, D.; Bajic, G.; Harrison, S. C.; Wesemann, D. R. Memory B Cell Repertoire for Recognition of Evolving SARS-CoV-2 Spike. bioRxiv Prepr. Serv. Biol. 2021. https://doi.org/10.1101/2021.03.10.434840.

(18) Schneider, M. M.; Scheidt, T.; Priddey, A. J.; Xu, C. K.; Hu, M.; Devenish, S. R. A.; Meisl, G.; Dobson, C. M.; Kosmoliaptsis, V.; Knowles, T. P. J. Microfluidic Antibody Affinity Profiling for In-Solution Characterisation of Alloantibody - HLA Interactions in Human Serum. bioRxiv Prepr. Serv. Biol. 2020, 2020.09.14.296442. https://doi.org/10.1101/2020.09.14.296442.

(19) Schneider, M. M.; Emmenegger, M.; Xu, C. K.; Condado Morales, I.; Turelli, P.; Zimmermann, M. R.; Frey, B. M.; Fiedler, S.; Denninger, V.; Meisl, G.; Kosmoliaptsis, V.; Fiegler, H.; Trono, D.; Knowles, T. P. J.; Aguzzi, A. A. A. Microfluidic Affinity Profiling Reveals a Broad Range of Target Affinities for Anti-SARS-CoV-2 Antibodies in Plasma of Covid Survivors. medRxiv 2020, 2020.09.20.20196907. https://doi.org/10.1101/2020.09.20.20196907.

(20) Fiedler, S.; Piziorska, M. A.; Denninger, V.; Morgunov, A. S.; Ilsley, A.; Malik, A. Y.; Schneider, M. M.; Devenish, S. R. A.; Meisl, G.; Kosmoliaptsis, V.; Aguzzi, A.; Fiegler, H.; Knowles, T. P. J. Antibody Affinity Governs the Inhibition of SARS-CoV-2 Spike/ACE2 Binding in Patient Serum. ACS Infect. Dis. 2021. https://doi.org/10.1021/acsinfecdis.1c00047.

